# Multi-purpose controllable protein generation via prompted language models

**DOI:** 10.1101/2024.11.17.624051

**Authors:** Zeyuan Wang, Binbin Chen, Keyan Ding, Jiawen Cao, Ming Qin, Yadan Niu, Xiang Zhuang, Xiaotong Li, Kehua Feng, Tong Xu, Ningyu Zhang, Haoran Yu, Qiang Zhang, Huajun Chen

## Abstract

Deep learning is increasingly powerful for designing proteins that meet structural and functional requirements. However, most existing methods follow a conventional pipeline: first defining a backbone structure and then generating sequences consistent with it. This approach, which encodes all design goals indirectly through structures, restricts flexibility and struggles to address multiple, complex design objectives simultaneously. We present PROPEND, a multi-purpose protein sequence design method based on the “pre-train and prompt” framework. We show PROPEND’s broad utility and accuracy both *in silico* and *in vitro* by directly controlling multiple properties through the prompt of backbones, blueprints, functional tags, and their combinations. For the five sequences tested with *in vitro* experiments, PROPEND achieved a maximum functional recovery of 105.2%, significantly outperforming the classical design pipeline’s 50.8%.

## Introduction

Designing novel proteins with specific, desirable properties, a process known as *de novo* protein design, is of great importance in synthetic biology and drug discovery. Traditionally, this field has relied on physics-based methods, which use optimization algorithms to search for sequences that correspond to the lowest energy configurations within a sequence–structure landscape. However, the vast number of potential rotamer combinations at different positions leads to a combinatorial explosion, posing considerable computational challenges (*1, 2*). Recently, the field has shifted toward AI-driven approaches that use conditional generative models trained on protein data, allowing for more efficient exploration of the sequence design space (*3–9*). This advancement points to a promising direction for developing proteins with optimized structural and functional properties.

Current AI-based methods typically adhere to the traditional protein design pipeline (*3–7*), which first defines a backbone structure and then leverages conditional generative models to identify sequences consistent with that structure (*10*). In this workflow, all design objectives, such as structural considerations and functional expectations, are indirectly encoded within the backbone itself, often through the spatial arrangement of active sites in the protein scaffold. However, capturing diverse design requirements solely through backbone structures can be limiting. Amino acids essential for maintaining structural stability may not overlap with those needed to meet specific functional goals. Precision is critical in achieving functional outcomes, as even small misalignments in the active site can compromise functionality (*11*).

In contrast, generating sequences directly based on design objectives presents a more effective approach (*8, 12, 13*) (Fig. 1A). Existing methods, however, often focus on a single design goal, limiting their ability to account for the diverse and complex requirements of protein functionality. This singular focus restricts their capability to optimize multiple properties simultaneously, which is crucial for advanced protein engineering. A framework capable of managing multiple design targets concurrently has the potential to transform the conventional protein design pipeline. By bypassing intermediary structural assumptions and focusing on direct sequence generation, such a strategy could result in more precise, adaptable, and efficient protein designs. The primary challenge lies in equipping protein generative models to fully understand and incorporate multiple design objectives within an integrated system.

**Figure 1:**
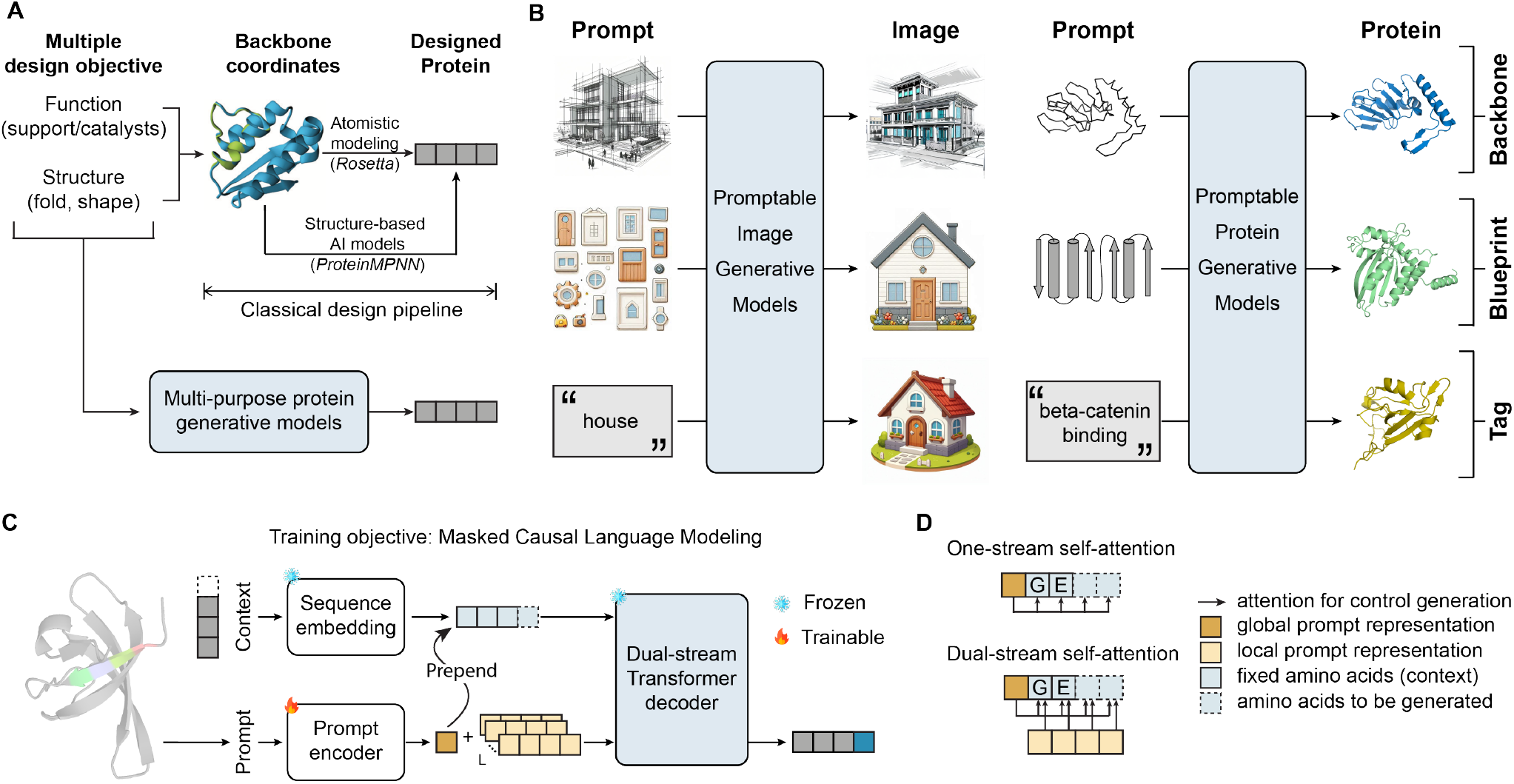
Multi-purpose controllable protein generation with prompted language models. (**A**) Most existing methods focus on single-purpose objectives, while the multi-purpose approach can handle multiple design scenarios simultaneously, offering a flexible and comprehensive solution for complex protein designs by balancing multiple constraints. (**B**) Prompting techniques are widely used to guide models in generating expected outputs. Similar to image generation, prompts in protein design can be categorized into backbone prompts, blueprint prompts, and tag prompts, each serving to direct the model toward specific design goals. (**C**) PROPEND uses global-local prompts to control the generative model. The global prompt defines the overarching design scenario (e.g., structure-based design), while local prompts provide detailed, context-specific instructions for each amino acid generated. (**D**) The traditional single-stream self-attention applies the same prompt for all amino acids, while dual-stream self-attention offers unique, detailed prompts for each amino acid during generation.

In recent years, the “Pre-train and Prompt” paradigm has gained widespread recognition in natural language processing and computer vision (*14–19*). Instead of training a model for each specific task, this approach utilizes crafted prompts to guide a generative model to produce desired outcomes. For example, prompts in the form of sketches, component layouts, or textual descriptions (Fig. 1B) enable models to generate a wide range of images (*20, 21*). Drawing inspiration from this approach, we investigate the use of prompt-based strategies for protein design, specifically targeting prompts based on protein backbones, secondary structure blueprints, and functional tags. This led to the development of PROPEND (PROmpt-guided ProtEiN Design), a unified framework that applies prompt engineering to enable multi-purpose applications of a single pre-trained protein language model (Fig. 1C).

We validate the effectiveness of PROPEND across various protein generation tasks. For protein inverse folding, PROPEND achieves a higher overall native sequence recovery rate (63.21% vs. 52.91%) and lower perplexity (3.13 vs. 4.02). PROPEND also generates proteins consistent with specified secondary structures and functional tags. Notably, in multi-objective sequence design, PROPEND designed a protein that folds according to the given structure and achieves a maximum functional recovery of 105.2%, significantly outperforming the classical design pipeline’s 50.8%, offering a robust and efficient alternative to traditional protein design approaches.

## Results

### Overview of PROPEND

PROPEND leverages prompts to unlock the potential of pre-trained protein language models (PLMs), enabling them to generate coherent and contextually relevant protein sequences across various design purposes. Given a generation purpose (e.g., a tertiary backbone structure), we first translate it into understandable prompts for the PLM, which then instruct the PLM to generate a protein sequence that satisfies the specified requirements.

The core of PROPEND consists of two types of prompts: global and local (Fig. 1C). Global prompts define the overall design context, such as specifying a tertiary backbone structure, providing broad guidance to the model. On the other hand, local prompts are condition-specific and offer detailed instructions about the position of each amino acid and its surrounding spatial information. These prompts are processed using a dual-stream self-attention mechanism, which allows each amino acid to have its own unique context (Fig. 1D). In this setup, the representation of each amino acid is updated by incorporating the preceding tokens along with both the global prompt and its associated condition-specific local prompt. This design enables precise control over various functional properties such as binding, catalysis, or structural support, and can even influence the local conformation of the amino acids being generated.

To further enhance controllable generation, PROPEND uses masked causal language modeling as its training objective. This approach, combined with a random masking strategy, introduces task complexity by adding uncertainty, which emphasizes the role of prompts in directing sequence generation. Additionally, PROPEND moves beyond the constraints of the language model’s discrete token space by embedding prompts in the latent space using a neural network and optimizing them through gradient descent. This allows for more flexible and effective prompt tuning.

### Generating protein sequence with the prompt of backbones

We first evaluated the effectiveness of PROPEND in solving the classical inverse protein-folding problem, where the atomic-level structure of a protein backbone serves as a prompt to guide the sequences generation. To benchmark PROPEND, we compared it against three state-of-the-art methods: ProteinMPNN (*22*), ESM-IF (*4*), and PiFold (*23*). This evaluation used the CATH v4.2 dataset and its standard splits (*3*), as defined in prior studies, to ensure consistency and comparability in results.

Our evaluation assessed both sequence accuracy and prediction certainty, measured by sequence recovery and model perplexity, respectively. PROPEND achieved a higher native sequence recovery rate than the best-performing baseline (63.21% compared to 52.91%) and a lower perplexity (3.13 vs. 4.02) (Fig. 2A). We further analyzed sequence recovery across different secondary structures (‘H’ stands helix, ‘E’ for sheet, and ‘C’ for coil) and varying levels of burial (reported by the average C*β* distance of the eight closest neighbors). PROPEND demonstrated consistent performance improvements across all spatial conformations, with gains most notable in surface-exposed regions (Fig. 2B). This suggests that PROPEND can generate plausible sequences even in the absence of explicit structural information. Given that multiple sequences can fold into similar structures, achieving 100% native sequence recovery is unattainable for protein design models. However, PROPEND showed excellent calibration, as evidenced by the resemblance between the substitution matrix of our method and that of BLOSUM62 (Fig. 2C).

**Figure 2:**
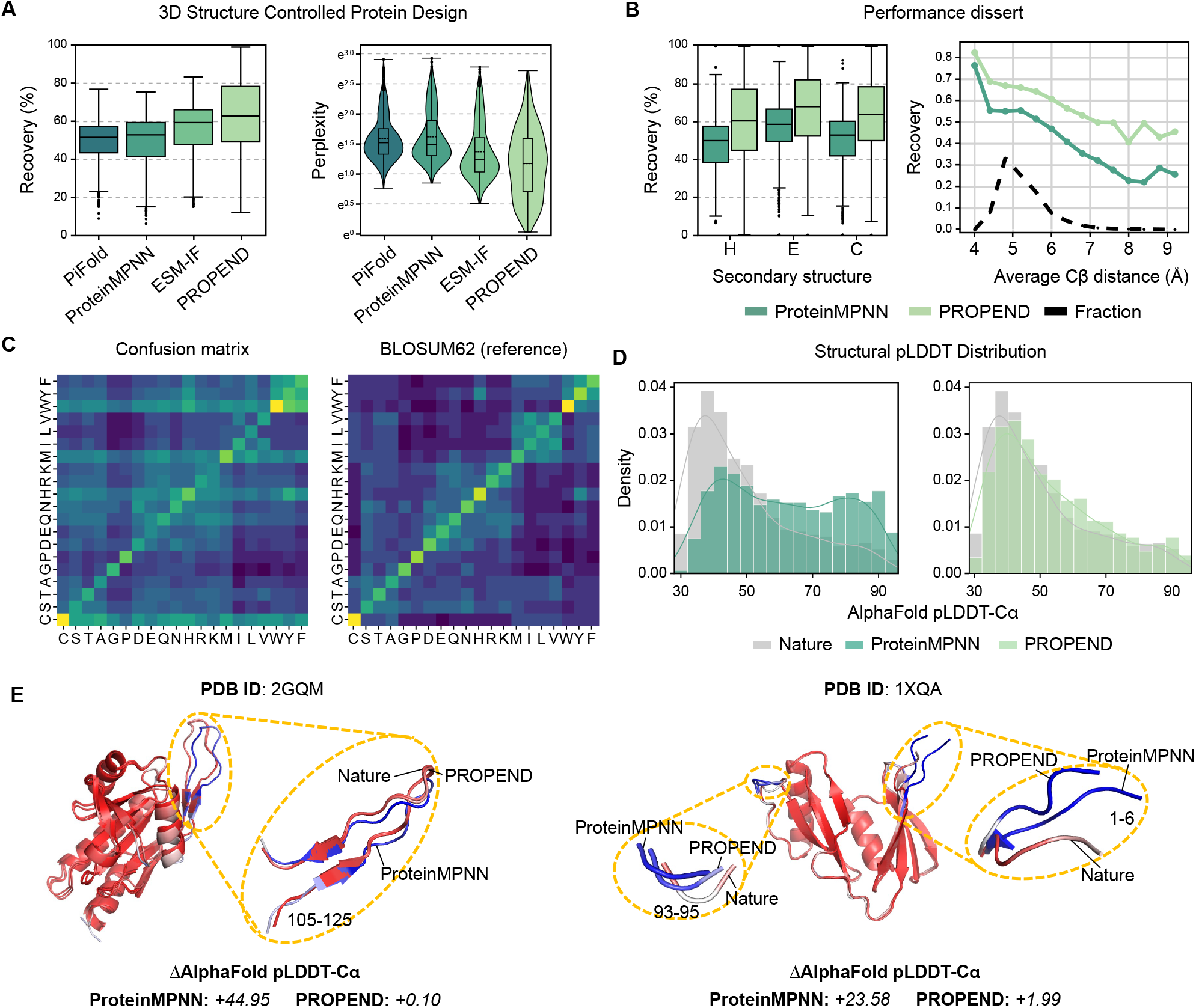
PROPEND faithfully follows the 3D structure prompts to generate protein sequences. (**A**) PROPEND has higher native sequence recovery and lower perplexity on CATH dataset (n=1120 structures). (**B**) Sequence recovery breakdown to residues on different secondary structures and burial. PROPEND outperforms other models at all kinds of secondary structures and all levels of burial. (**C**) Confusion matrix of PROPEND, compared to BLOSUM62 as reference. (**D**) Distribution of sequence pLDDT from the test set (grey), ProteinMPNN (dark green) and PROPEND (light green). (**E**) The predicted structures of sequences designed by ProteinMPNN and PROPEND for two structures (2GQM and 1XQA), compared to the native ones.

To assess foldability, we used AlphaFold2 (*24*) to predict local distance difference test (pLDDT) scores for PROPEND-generated sequences compared to those generated by ProteinMPNN, with native sequences as references. While pLDDT scores above 70 are considered indicative of high prediction confidence, lower pLDDT scores can be consistent with intrinsically disordered regions (IDRs) of proteins (*25*). PROPEND-generated sequences yielded pLDDT distributions that closely mirrored those of native sequences (Fig. 2D). Detailed analysis of representative examples (PDB ID: 2GQM and 1XQA) with different pLDDT scores from PROPEND and ProteinMPNN illustrates that PROPEND not only maintains high-confidence regions but also accurately reflects the low-confidence regions (Fig. 2E). This suggests that PROPEND is better at preserving the native-like distribution of structural confidence across the entire protein sequence compared to ProteinMPNN. Such fidelity in maintaining the structural characteristics of native proteins highlights PROPEND’s potential in generating biologically relevant protein sequences.

### Generating protein sequences with the prompt of blueprints

Next, we investigate whether PROPEND can generate protein sequences with the desired properties using less informative prompts than backbones, such as blueprints. Classical approaches in protein design frequently rely on simplified blueprints of protein folding topologies, defined as the identity and sequence of secondary structure elements. While these blueprints are less detailed than 3D backbones, they offer greater flexibility, making it possible to design novel fold topologies that do not exist in nature. This makes them particularly valuable in protein design research (*26*).

To evaluate this capability, we curated a dataset of non-redundant secondary structures (*27*) as prompts and generated protein sequences directly, bypassing the intermediate step of assembling the protein backbone. Because the prompt of blueprints provides a broader design flexibility than the backbone prompt, we assessed the design accuracy by predicting the 3D structures of generated sequences and then applied the Dictionary of Secondary Structure in Proteins (DSSP) algorithm to compute the aligned residue-wise sequence similarity (*28*). Over the generations using secondary structure in test set, we observed that both *de novo* designed and inpainted sequences generated by PROPEND closely aligned with the secondary structure distributions of natural sequences including random coils and well-defined structural elements (Fig. 3A), while maintaining low sequence similarity (Fig. 3B). Considering the constraints imposed by the physical and chemical properties of amino acids, the alignment between coarse-grained secondary structures and fine-grained structural details is crucial. The 8-state secondary structure confusion matrix has confirmed that PROPEND understands protein design blueprints and generates candidate proteins that closely align with the intended design (Fig. 3C). To evaluate whether PROPEND could design novel and diverse sequences from the broader design space, we examined 200 designed sequences under the same blueprint. The result of high recovery coupled with low similarity suggest that our blueprint prompts effectively guide PROPEND to the appropriate protein design subspace (Fig. 3D).

**Figure 3:**
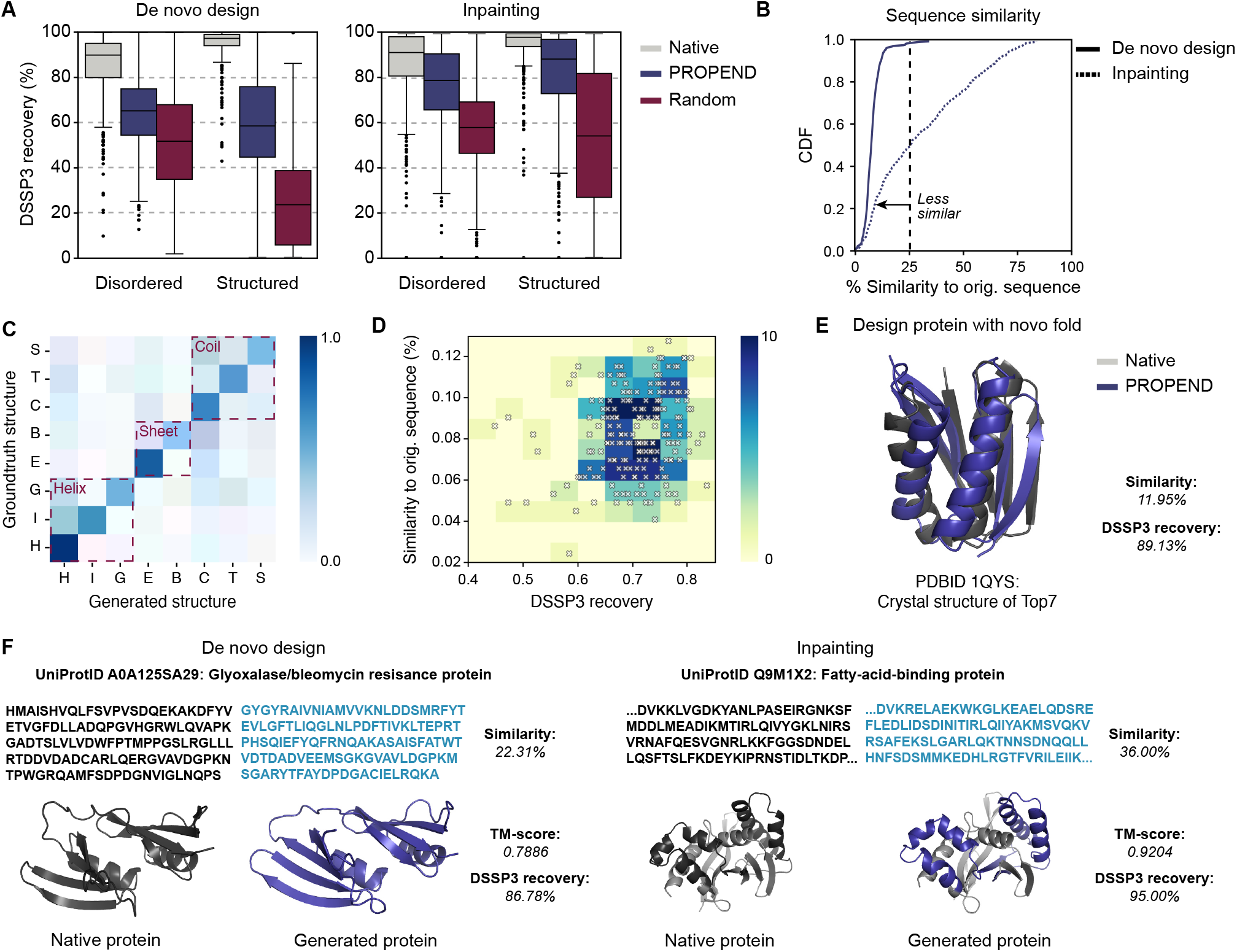
PROPEND generates sequences with “blueprint” (identity and order of secondary structure elements). (**A**) Distributions of secondary structure recovery scores for de novo designed and inpainted sequences. (**B**) Distribution of sequence similarity relative to the original protein for generated proteins from de novo designed and inpainted sequences. (**C**) Secondary structure confusion matrix of PROPEND. (**D**) DSSP3 recovery versus sequence similarity for sequences under the same secondary structure (n=200). (**E**) Predicted structures and metrics for Top7, a protein that folds into stable new topologies not seen in nature, and generations from PROPEND. (**F**) Generated sequences, predicted structures, and computed metrics for representative design examples from PROPEND. The original sequence is shown in black and the generated sequence is highlighted in blue.

Designing topological structures that are not seen in nature could have significant implications in materials biology and medicine. This is a particularly challenging task for traditional MSA-based models since there are no reference sequences for such designs. We are curious whether prompt-based methods can overcome this obstacle. We choose the Top7 protein (*26*), a 93-residue *α*/*β* protein, as an example. Compared with traditional optimization algorithms, PROPEND offers advantages in reduced computation and faster design speed, successfully delivering sequences that meet specified conditions (Fig. 3E). To further demonstrate these capabilities, we visualize predicted structures (Fig. 3F) for a sample of high pLDDT generations controlled by the blueprint prompts that exhibit low sequence similarity and high secondary structure recovery. Together, these results confirm that PROPEND can robustly generate diverse proteins guided by blueprint prompts, showcasing its potential to design novel structures beyond the limitations of conventional models.

### Generating protein sequences with the prompt of functional tags

Traditional protein design methods define “function” by embedding an active site into a specified protein “scaffold”. However, function and structure are distinct attributes; proteins with similar structures can have vastly different functions due to unique dynamics (*29*). Here, we investigate the potential of using functional tags directly as prompts to guide protein design.

Using functional tags from UniProtKB (*30*), our generation results span a broad landscape of natural protein sequences (Fig. 4A), representing seven functional categories with diverse folds, active site architectures, and enzymatic mechanisms. To assess the functional plausibility of these generated sequences, we utilize foldseek (*31*) to search for proteins in PDB (*32*) that exhibit a structure most similar to the predicted structure. Results show that the identified proteins all possess functions consistent with the provided functional tags(Fig. 4A). We further evaluated sequence diversity by calculating the residue-wise sequence similarity between generated sequences and their closest counterparts in the training dataset, finding PROPEND-generated sequences to have low similarity to known functional proteins (Fig. S22). Notably, PROPEND outperformed a state-of-the-art tag-controlled language model, ProGen, both qualitatively and quantitatively (Fig. S33).

**Figure 4:**
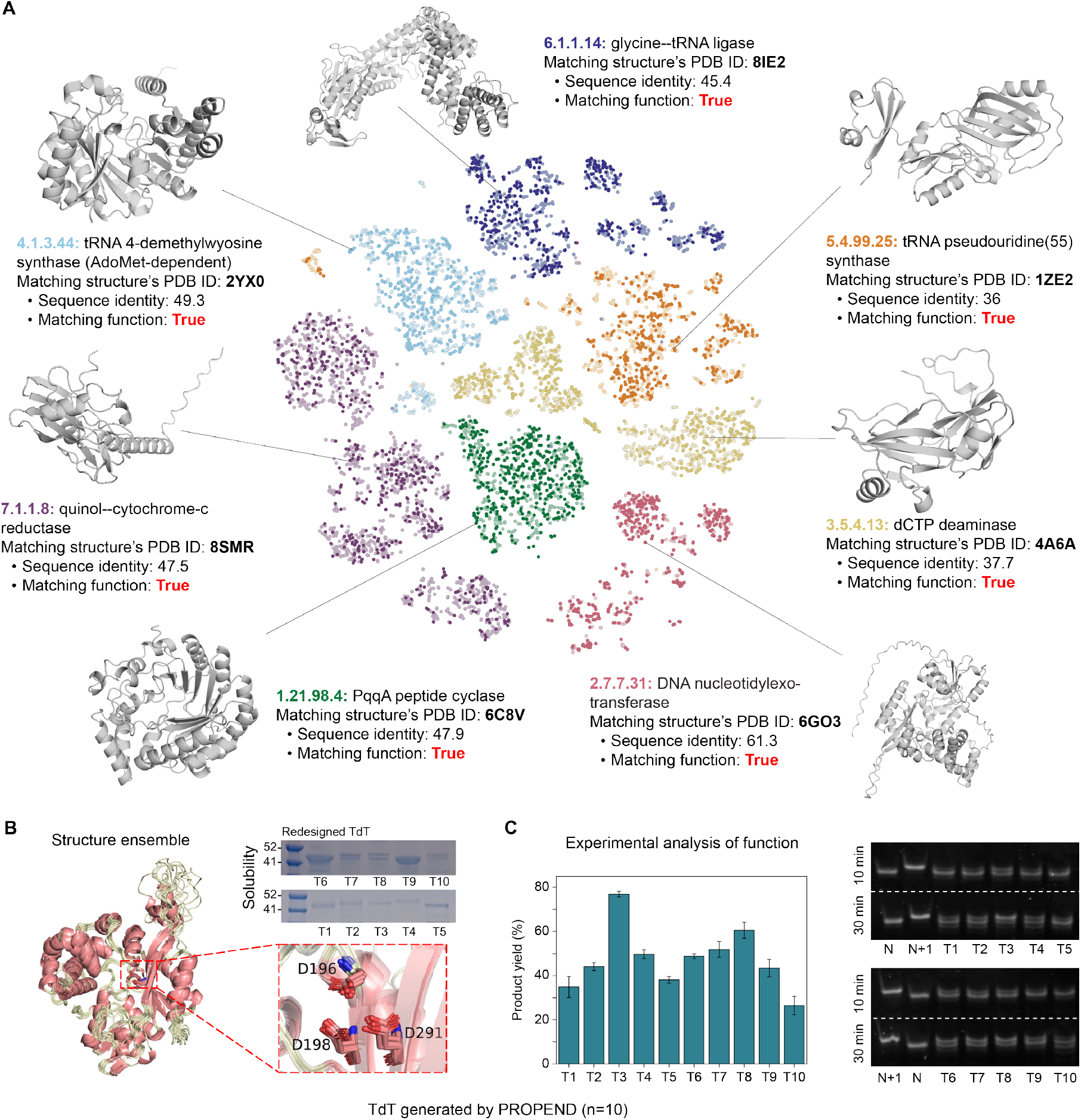
PROPEND enables understanding natural language and generating corresponding proteins. (**A**) t-SNE of ESM2 embeddings of natural sequences (light color) and generated sequences (dark color) from PROPEND using protein functions as prompts. Predicted structures and metrics for representative functional plausible generations from PROPEND. (**B**) The predicted structure alignment and expression of ten sequences designed by PROPEND using functional tag-DNA nucleotidylexotransferasem-as prompt, showing conserved catalytic residues. (**C**) The activity of TdTs detected by Urea-PAGE for primer extension reactions. The primer extension reaction was carried out at 37 oC for 10 min and 30 min. N, 23 nt primer control; N+1, 24 nt product control.

Besides the *in silico* validation of PROPEND, we also conducted a *in vitro* experiment to evaluate its performance in addressing Terminal Deoxynucleotidyl Transferase (TdT) design challenges, a template-independent DNA polymerase for *de novo* DNA synthesis. With the functional tag “DNA nucleotidylexotransferase” as a prompt, PROPEND generated 1,000 sequences with lengths varied from 120 to 998. We refined this library by excluding sequences shorter than 300 amino acids, those with significantly low sequence identity to wild-type sequences, and sequences lacking the three critical catalytic aspartic acids (Fig. S4A). From this refined set, we selected ten sequences exhibiting 40-60% identity to natural TdT for *in vitro* evaluation. Each sequence expressed solubly in *Escherichia coli* and purified using NiNTA (Fig. 4B). We conducted the template-free nucleotide addition reaction with a 23 bt primer and 2’,3’-dideoxyguanosine-5’-triphosphate (ddGTP) as the substrate. The reaction products were detected by the polyacrylamide gel electrophoresis. All sequences demonstrated detectable activity after 30 minutes, with S3 exhibiting higher activity than others (Fig. 4C). These results demonstrate that PROPEND can use complex natural language as cues to generate novel, diverse, and structurally plausible functional proteins.

### Generating protein sequences with multi-purpose prompts

Achieving multiple property requirements simultaneously is a practical yet challenging goal in protein design. Given PROPEND’s capacity for multi-purpose generation, we hypothesized that it could effectively integrate multiple prompts to generate sequences meeting complex criteria.

Firstly, our focus was on designing multi-domain proteins by leveraging PROPEND’s dual-stream self-attention mechanism to precisely establish positional relationships among domains within a sequence (Fig. 5A). To test this, we generated 200 sequences with fixed scaffold regions, supplying two domain tags (SH2 and PTB/PI domains) as conditioning information. PROPEND’s inpainting of these domains resulted in PLM representation distributions that closely matched those of natural protein sequences (Fig. 5B). Structural analysis confirmed that the generated domains maintained isolation while preserving structural integrity (Fig. 5C), indicating that PROPEND effectively supports multi-domain design through multifunctional prompts.

**Figure 5:**
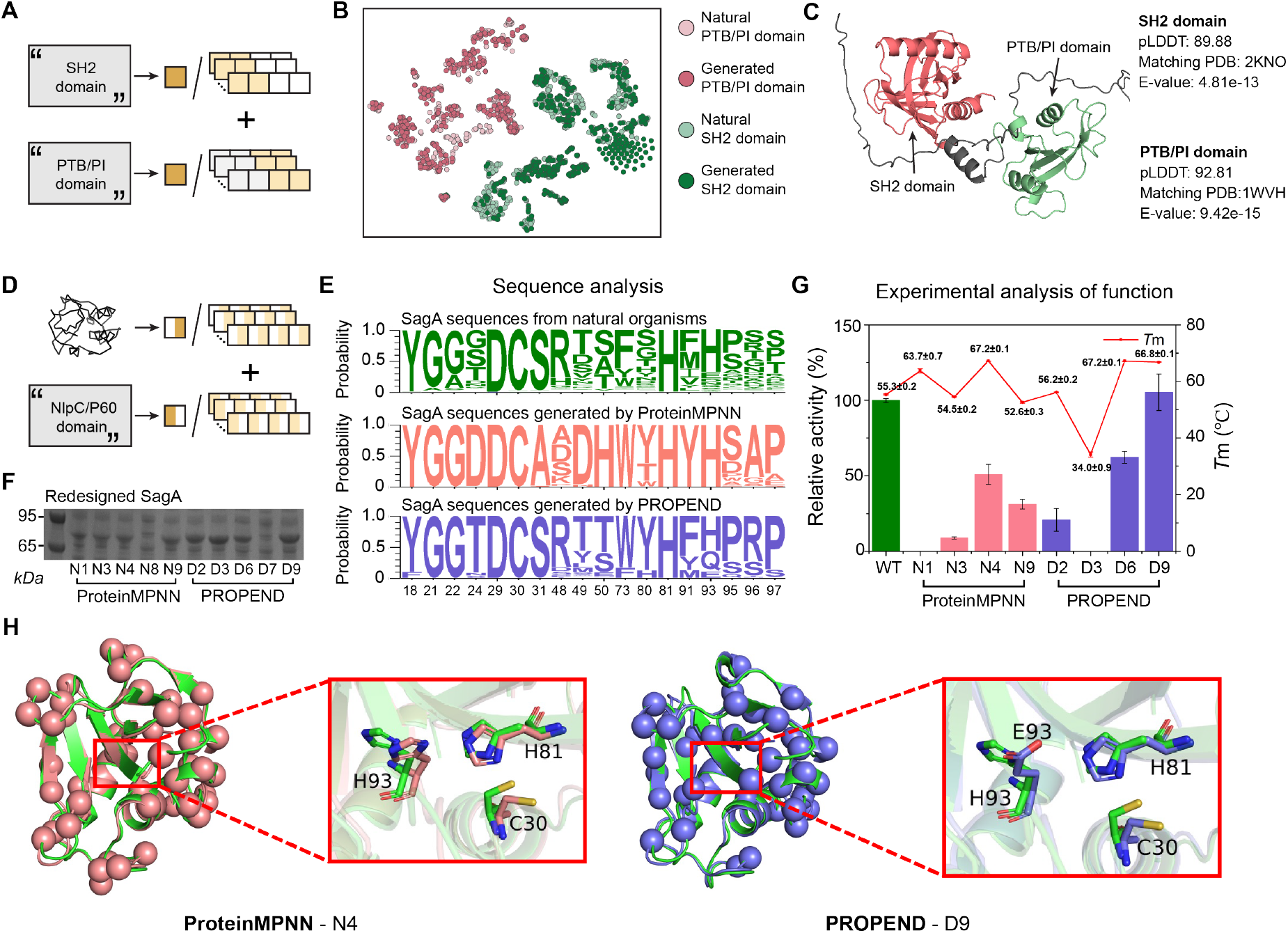
PROPEND takes multiple design objectives into consideration simultaneously. (**A**) Multi-domain sequences are generated from PROPEND using concatenating multiple subsequence domain prompts. (**B**) t-SNE visualization of ESM2 embeddings for domains within multi-domain proteins, generated using the PROPEND framework under multiple functional prompts. (**C**) Predicted structures and metrics for representative generations. (**D**) PROPEND considers both functional and structural design objectives, achieved through the summation of the corresponding prompts for each. (**E**) Expression of generated SagAs. (**F**) Sequence alignment of SagAs from natural organisms (Green), designed by ProteinMPNN (Red) and PROPEND (Blue). ProteinMPNN uses only structure as a condition, while PROPEND directly employs both structure and functional tags as prompts. (**G**) Activity and thermostability of SagAs generated by ProteinMPNN and PROPEND. (**H**) Structures of two active SagA variants N4 and D9. Green, wild type; Cyan and blue, SagAs generated.

Next, we redesigned the NlpC/p60 peptidoglycan hydrolase secreted antigen A (SagA), focusing on both its structural and functional characteristics. For comparison, we generated sequences with ProteinMPNN using SagA’s functional backbone structure from the RCSB PDB (ID: 6B8C), while PROPEND utilized this structure with additional NlpC/P60 functional tags for controllable generation (Fig. 5D).

Functional site conservation was examined by analyzing amino acid frequencies within 3 Åof the ligand, visualized using WebLogo3 (Fig. Fig. 5F). Key catalytic residues—Y18, D29, C30, S31, and H81—were highly conserved, while variability emerged at other sites across 23,779 sequences. A notable divergence was observed at position 31, where sequences generated by ProteinMPNN contained alanine, contrasting with the serine found in both natural sequences and those generated by PROPEND. Additionally, outside these five conserved residues, ProteinMPNN introduced sequence conservation at positions 24, 50, 73, 91, 93, 95, and 96, diverging from both natural sequences and PROPEND’s output (Fig. 5E). This suggests PROPEND provides broader diversity in SagA’s catalytic region, aligning closely with natural variability.

We selected the top 10 lowest-scoring sequences from PROPEND and ProteinMPNN and expressed five from each with low sequence identity in *E. coli*: N1, N3, N4, N8, N9 (ProteinMPNN) and D2, D3, D6, D7, D9 (PROPEND). We obtained the 3D structures of these artificial SagA sequences by AlphaFold2, and aligned them with the X-ray crystal structure of SagA (PDB ID:6B8C). We found they had the same secondary structure, including *α*-helix, *β*-folding and loop structure at the same position, confirmed that both methods accurately preserved SagA’s secondary structure features. All selected sequences were solublly expressed in *E. coli*, except for N8 from Protein-MPNN and D7 from PROPEND (Fig. 5F). Notably, the sequences generated by PROPEND exhibited higher expression levels compared to those from ProteinMPNN. We also measured the activities and thermostabilities of these generated sequences (Fig. 5G). While most sequences demonstrated lower activities than the wild-type SagA, D9 from PROPEND displayed activity comparable to the wild type. Additionally, melting temperature analysis revealed that sequences N1, N4, D6, and D9 exhibited enhanced thermostability compared to the wild type, highlighting the potential of PROPEND-generated proteins in maintaining structural integrity under varying thermal conditions. Structure analysis revealed a mutation H93E in the active center of D9 (Fig. 5H). Although histidine is the most frequent amino acid found in natural sequences, a previous study showed mutation at this position has the potential to increase the activity of SagA (*33*).

## Discussion

PROPEND represents one of the pioneering attempts to use AI models for generating protein sequences directly conditioned on diverse design objectives. This approach is particularly effective when addressing design goals that extend beyond tertiary structures, where traditional design pipelines often encounter limitations. PROPEND achieves this by leveraging a highly efficient prompt-injection architecture, enabling multi-purpose applications from a single pre-trained protein language model. Since the generation of amino acids in language models depends on the provided context, and the model has been trained on universal sequences, finding the appropriate prompts allows for efficient exploration of sequence search spaces corresponding to the design objectives.

From a practical perspective, PROPEND’s efficacy in tackling various design challenges demonstrates that prompt engineering serves as an effective modular controller for guiding general pretrained language models without necessitating modifications to their trainable parameters. This flexibility allows for the simultaneous pursuit of multiple design goals. In contrast to traditional pipelines, which typically construct tertiary structures for multiple objectives before designing corresponding sequences, PROPEND directly integrates these design goals, resulting in more efficient modeling of conserved patterns. This efficiency is particularly evident in tasks like the design of TdT using approaches such as ProteinMPNN.

We view PROPEND as a foundational step toward developing AI methods capable of end-to-end protein sequence design with integrated desired properties. To advance this foundation, two key improvements can be explored. First, moving beyond the left-to-right autoregressive language model could enhance design flexibility. By enabling the model to reason over arbitrary pairwise dependencies and incorporating inductive priors based on conservation patterns, PROPEND’s predictive capabilities can be further refined. Second, while PROPEND currently relies on a data-driven likelihood distribution, embedding fundamental physical and chemical principles within the model could expand its design scope. This integration could improve its performance in data-scarce areas, such as non-proteinogenic amino acids, and potentially extend its design capabilities beyond residues and supporting all-atom biomolecular design.

In summary, we present a prompt-guided protein design framework that breaks away from the constraints of traditional design pipelines. By leveraging the data-driven and expressive power of protein language models, PROPEND offers a new approach to protein design. It can be directly applied to structure-based, function-based, or multi-objective-based generation of protein sequences, and it may be extended to accommodate more specific experimental needs, such as pH, temperature, or functional participants. We envision that PROPEND, through the creation of a system of design objectives translated into modular prompts, will enable precise *de novo* protein design to address challenges in biology, medicine, and environmental science.

## Author contributions

Z.W. and K.D. conceived the study and designed the methods. Z.W. and B.C. implemented and conducted the experiments. Z.W., B.C., K.D., and J.C performed the analyses. Z.W., B.C., K.D., J.C., H.Y., Q.Z., and H.C. provided suggestions on the experimental analysis. Z.W., B.C., K.D., M.Q., X.Z., X.L., H.Y., Q.Z., H.C. wrote and revised the manuscript. Y.N., K.F., T.X., and N.Z. provided suggestions on the model design and writing. H.Y., Q.Z., and H.C. supervised the project. All authors have read and approved the manuscript.

## Competing interests: The authors declare no competing interests

## Data and materials availability

The foundation model used ProGen2 from https://github.com/salesforce/progen/. The backbone prompt encoder used ProteinMPNN v_48_010 from https://github.com/dauparas/ProteinMPNN. The blueprint and functional tag prompt encoder used Chemical Language Model from https://huggingface.co/GT4SD/multitask-text-and-chemistry-t5-small-augm. The dataset for training is available at https://doi.org/10.5281/zenodo.14006102.

## Supplementary materials

Materials and Methods

Figs. S1 to S5

Tables S1

References *(33-40)*

## Supplementary Materials

## Materials and Methods

### Datasets

The datasets used to tune modular prompts fall into three main categories based on their design objectives: tertiary structure, secondary structure, and functional annotation. In this subsection, we describe the curation of each dataset.

### Tertiary structural datasets

We used the CATH v4.2 dataset, a well-established benchmark for protein structure analysis (*3*). This dataset contains 19,700 high-resolution single-chain structures sourced from the Protein Data Bank (PDB) (*32*). These structures were split into training, validation, and test sets with sizes of 18,024, 608, and 1,120, respectively. The division was based on the CATH v4.2 40% non-redundant protein set.

### Secondary structural datasets

We used the PS4 dataset (*27*), one of the largest secondary structure datasets, built from the full pre-computed DSSP database (*28*). The dataset was filtered to 40% sequence similarity using Levenshtein distance, yielding 18,731 non-redundant protein chains. These were further split into 17,799 training samples and 932 validation samples.

### Functional datasets

We compiled a comprehensive protein sequence dataset by integrating all entries from UniProtKB/Swiss-Prot and those from UniProtKB/TrEMBL with an annotation score above 3, resulting in 14.14 million non-redundant sequences (*30*). The functional annotations include molecular functions (from Gene Ontology), enzyme commission (EC) numbers, and domains, covering 28,790 hierarchical terms. The accepted natural language names of these functions were used as prompts to guide protein generation. Sequences without these annotations or with lengths shorter than 1,024 amino acids were excluded, yielding a final dataset of 13.45 million sequences. This dataset was then split into 12 million sequences for training and 1.45 million for validation. During prompt tuning, unaligned protein sequences were presented to the model, with a randomly selected non-empty subset of functions serving as instructions.

### The Proposed Model

#### Preliminary

Language models are a class of generative models that learn to generate data from context. They acquire this ability by modeling statistical relationships from text documents during a self-supervised training process. Let *x* = (*x*_1_, …, *x*_*n*_) be a sequence of tokens in corpus χ. Language modeling aims to maximize the probability distribution of *x*. This involves breaking down the probability of *x* into the probabilities of each individual token within the sequence, taking into account the context provided by the preceding tokens. The learned process is parameterized by a model such as a neural network *θ* with minimizing the negative log-likelihood over χ:

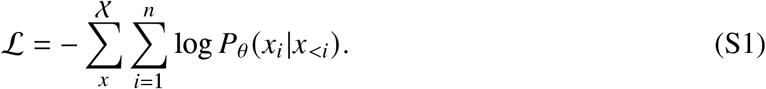

Pre-trained language models have demonstrated the ability to learn general-purpose representations from vast amounts of raw data. These models can be effectively adapted to various downstream tasks by using prompts. A prompt is a sequence of tokens *p* = (*p*_1_, …, *p*_*m*_) that describes the task at hand, such as ”write a poem.” When a prompt is prepended to the input, the model generates responses in an autoregressive manner, conditioned on both the prompt and the preceding context (*P*(*x*_*i*_) = *P*_*θ*_ (*x*_*i*_ |*x*_*<i*_, *p, θ*)). More concise prompts tend to elicit more focused and relevant responses from the model. The practice of prompt engineering, which involves designing and refining these prompts, has been developed across various modalities, including text, images, and audio, and has been further optimized by techniques such as chain of thought.

#### PROPEND

This work presents a PROpmt-guided ProtEiN Design (PROPEND) model that leverage (1) global-local prompt and masked causal language modeling and (2) dual-stream self-attention mechanism to design protein from tertiary structure, to secondary structure, to function-guided. Prompt engineering operates by creating a prompting function that results in the most effective performance on the downstream task. One must first consider the prompt shape, and then decide whether to take a manual or automated approach to create prompts of the desired shape. This section describes the prompts for multiple design objectives and architecture. A prompt typically consists of both an instruction and relevant context. For example, when extracting specific details from a text, one might begin with an instruction like, ‘Identify the names of places in the following passage,’ followed by the text itself. Similarly, in protein generation, we designed global and local prompts for analogous purposes. The global prompt *p*^*G*^ specifies the overall design scenario, while the local prompt *p*^*L*^ provides detailed descriptions of the design content. Since the more descriptive and detailed the prompt is, the better the results. In the local prompt, we use the residue-level description to guide the generation. Then the proteins generated under the guidance of the prompt can be formulated as

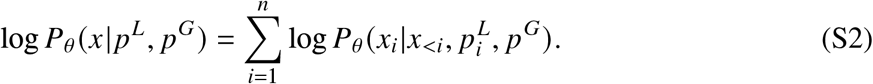

Instead of using a limited discrete vocabulary to search for prompts, which doesn’t fully align with the continuous nature of neural networks, we generate prompts directly in the latent space. Specifically, given a design objective—such as a protein backbone structure—we utilize a prompt encoder to embed the objective as 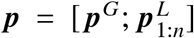. We then optimize the encoder’s parameters through gradient descent to identify the most effective prompt.

Well-established PLMs excel in sequence completion. This means that, given prior context, they can generate plausible continuations based on pre-learned knowledge without relying on prompts. However, this capability can hinder prompt search. To mitigate this, we propose masked causal language modeling (MCLM), which introduces slight perturbations to the input sequence, ensuring the model remains dependent on the prompt for more effective optimization. In this approach, a fraction of amino acids in each sequence are masked, introducing controlled corruption. The network is trained to generate informative prompts that guide PLM output:

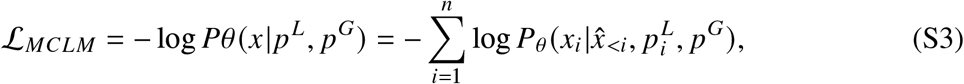

where 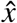 represents the corrupted context, generated by sampling a set of indices *M* for each sequence *x* and replacing the true token at each index *i* with a mask token.

#### Model architecture

PROPEND is parameterized with multiple prompt encoders tailored for diverse design objectives, an adaptor, and a modified Transformer decoder (Fig. S1). We utilize message-passing neural networks to encode protein 3D structures and Transformer encoders for discrete data like protein secondary structures and natural language 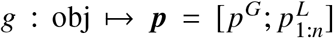 where obj is the design objective, *p*^*G*^ ∈ ℝ^*d*^ and 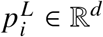, *d* is the hidden dimension. Due to the lack of a one-to-one correspondence between functional descriptions and amino acids, the local prompt associated with each amino acid remains consistent throughout the generation process, encapsulating all functional description information.

The global prompt is introduced before the protein sequence, enabling each generated amino acid to assess its impact via the self-attention mechanism. For the local prompt, the adapter is used to customize it for each layer, guiding protein generation through a modified Transformer decoder. Specifically, the modified Transformer decoder architecture comprises a series of decoder blocks that alternate between two functions: dual-stream multi-head self-attention (DSA), which computes the interaction of amino acids with their context, and a feedforward network applied independently to each amino acid. In the *l*-th layer, the input comprises a protein representation *h*^*l*−1^ ∈ ℝ^*n*×*d*^ from the (*l* − 1)-th layer and the corresponding local prompt *p*^*l*^ ∈ ℝ^*n*×*d*^, producing a sequence of updated protein representations ***h***^*l*^.

DSA module update each amino acid representation by computing a weighted sum of values of its preceding residual and the corresponding local prompt, with weights determined by a compatibility function between the query and the corresponding key. Specifically, we first generates a key, value, and query based on ***h***, and key and query of ***p***:

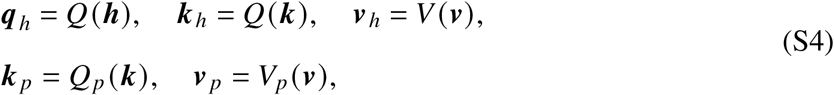

where ***q***_*h*_, ***k*** _*h*_, ***v***_*h*_, ***k*** _*p*_, and ***v*** _*p*_ are linear projections with dimension *d*_*k*_. Then an attention score between amino acids and prompt representations is calculated based the similarity between query to key:

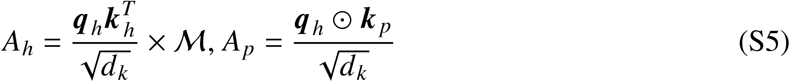

where M is a causal mask to prevent information leakage, is element-wise product and 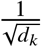 is a scaling factor. The output vectors at the attention head are the weighted average of their context value vectors:

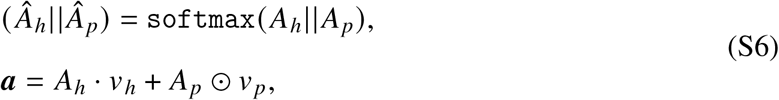

where || denotes concatenation. One step of self-attention directly models possible pairwise interaction between all amino acids and corresponding prompts in dual sequences simultaneously. Multi-headed self-attention concatenates the output of *t* independent attention heads ***a***_DMA_ = ***a***_1_ · · · ***a***_*t*_, which enables representation of different interposition interaction patterns. The feed-forward network is followed to map it to ***h***^*l*^ with dimension *d*. The final layer outputs are passed to a projection *W* to log probabilities ***y***.

#### Training details

The sequence decoder in PROPEND utilizes pre-trained ProGen2 models, which have demonstrated superior performance in protein language modeling. For prompt encoders, the ProteinMPNN architecture is employed to encode 3D backbone structures, while the T5 encoder is utilized to encode discrete secondary structure elements and functional descriptions. Since the tertiary/secondary structure describes the posture of each amino acid, we use the structural representation of each amino acid as its local prompt. There is no one-to-one correspondence between functional descriptions and amino acids. We introduce a special token [PROMPT] and prepend it in function description. The representation of this token is used as functional local prompt.

During prompt tuning, the sequence decoder remains frozen while the prompt encoders are optimized, resulting in only 10% of the parameters being trainable. Each encoder is trained from scratch using the AdamW optimizer, with a learning rate of 1e-4, a linear warmup over 2,000 steps, and dynamic batching to optimize GPU usage. ProGen2 models are available in various sizes; we primarily focus on the base and xlarge models, which have 764M and 6.4B parameters. 764M parameter models were trained on 4 24GB NVIDIA 4090 GPUs; 6.4B parameter models were trained on 4 32GB NVIDIA V100 GPUs. The maximum number of tokens per GPU in each batch was reduced from 8,192 to 1,024 to accommodate training the larger 6.4B parameter models. We trained each prompt encoder to minimize the negative log-likelihood, as defined in Eq. S3, on their respective datasets. The early stopping is used if there is no improvement on the perplexity on the validation set for 3 epochs. All models are implemented in PyTorch Lightning framework.

### Baseline models

To assess PROPEND’s performance across various design scenarios, we conducted an extensive comparison against established protein design models. The ProteinMPNN baseline (*22*) was trained using the same architecture and dataset as the PROPEND structural encoder. For a fair comparison, we retrained ESM-IF (*4*), originally trained on AlphaFold2-predicted structures and the CATH 4.3 dataset, using the CATH 4.2 dataset. ESM-IF often generated the end-of-sequence token (<eos>) prematurely, resulting in inconsistent lengths between backbone and generated sequence. To address this, we manually set the logits for <eos> to infinity and halted the generation when the sequence length matched the backbone length. PiFold (*23*) directly maps structure to amino acids; therefore, we excluded contextual information during perplexity calculation. In our function-based design baselines, we supplemented the control tags of ProGen (*8*) with previously missing functional descriptions. Following the fine-tuning of the model with sequences specific to each function, we utilized ProGen to generate artificial sequences by inputting the corresponding function description.

### In silico evaluation methods

#### Generating protein sequence with the prompt of backbones

Starting with a 3D-structure-based design prompt, new sequences were generated autoregressively, one amino acid at a time. Since there is a direct one-to-one relationship between the backbone and its sequence, the generation process concludes when the amino acid linked to the final local prompt is produced. The generated query sequence was evaluated relative to the corresponding original sequence using the perplexity and recovery. Perplexity quantifies the uncertainty of a sample drawn from a discrete probability distribution, calculated as the inverse likelihood of the native sequences ***x*** under the model’s predicted sequence distribution:

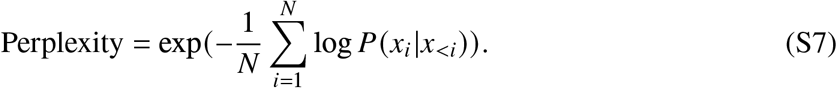

To maximize sequence recovery, the sequences are sampled using the argmax function from the model’s predictions. We also evaluated the substitution scores between native and sampled sequences using a log-odds ratio formula, analogous to the one employed in the BLOSUM62 matrix. For any two amino acids *a* and *b*, the substitution score *s*(*a, b*) is calculated as:

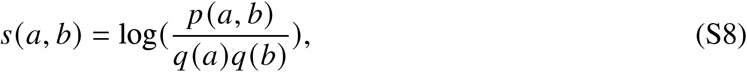

where *p* (*a, b*) denotes the jointly probability of substituting native amino acid *a* with sampled amino acid *b*, while *q* (*a*) and *q* (*b*) represent their respective marginal probabilities in the native and sampled distributions. Across generated sequences both the confusion matrix and its Pearson correlation coefficient with BLOSUM62 were reported. The generated sequences were further evaluated by comparing their predicted local distance difference test (pLDDT) distribution to that of their native counterparts. For each sequence, pLDDT and structures were predicted using AlphaFold without incorporating multiple sequence alignments or template information.

#### Generating protein sequences with the prompt of blueprints

Native sequences and their corresponding secondary structures were sampled from the PS4 dataset as input for PROPEND models. The eight secondary structure classes were consolidated into three major states: helix (including 3_10_ helix, *α* helix, and *π* helix), sheet (encompassing isolated *β*-bridge residues and strands), and coil (bends, turns, and coils). The native sequence served as the query, while the secondary structure functioned as the blueprint prompt. All sequences were subsampled to a length of 512 residues. For de novo design, PROPEND used the 3-state secondary structure as the prompt, guiding the generation of new residues. For protein inpainting, PROPEND was employed to design half-length continuous sequences. Since the sequence is generated autoregressively from left to right, the model used the native sequence preceding the missing segment as context. The blueprint prompt guided the generation of the missing region, after which the remaining half of the native sequence was manually attached. For sequence ensemble generation, we use temperature *T* = 1 and perform random sampling within the protein design space.

The generated sequences were input into ESMFold for 3D structure prediction. The per-residue secondary structures were then obtained using the DSSP algorithm. A single-sequence structure predictor (ESMFold) was used in place of MSA-based structure prediction methods, because of the observed degradation in prediction performance when dealing with de novo proteins lacking homologous information. Secondary structures for native sequences, generated sequences, and randomly generated sequences were computed to evaluate the performance of PROPEND predictions. The randomly-sampled baseline was constructed by randomly sampling amino acids across the region where design was required. A direct comparison between the original sequence and the generated sequence guided by secondary structure was conducted by calculating the sequence similarity between the fraction of shared residues and secondary structure elements between the two sequences. 3D structural similarity was measured via the template modeling score (TM-score) for the two predicted structures following structural alignment:

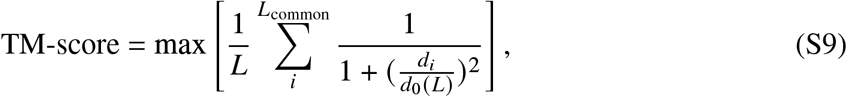

where *L* is the length of the secondary structure; *L*_common_ is the number of shared residues; *d*_*i*_ is the distance between the *i*^*th*^ pair of residues; and 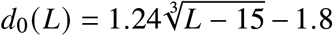 is a distance scale for normalization.

#### Generating protein sequences with the prompt of functional tags

In function-based design, we selected seven representative enzymes based on the chemical reactions they catalyze (oxidoreductases, transferases, hydrolases, lyases, lsomerases, ligases, and translocases). 200 samples were generated for each unique function. To generate a functional protein with PROPEND, the context was initialized to the global prompt representing function-based design and the function description is encoded into a local prompt of length 1. Each amino acid to be generated updates its representation based on the previously generated sequence and the global and local prompts. This process continuous until the <eos> token is produced, indicating that a complete protein has been generated. In this approach, on average, the functional proteins generated by PROPEND contained substantial sequence diversity with average sequence varying between 187-507 across families.

Previous work has shown that even without explicit supervision, protein language model embeddings contain information about both sequence and function as captured in GO annotations (*34,35*). To evaluate function coverage, ESM2 embeddings were computed for each of 200 generated protein sequences and all native functional sequences. The resulting embeddings were visualized in 2D via t-distributed Stochastic Neighbor Embedding (t-SNE) fit to the native data and with `init`=pac and `perplexity`=10. We calculated the embedding space Fréchet distance between a set of generated sequences and the test set (Fréchet Protein distance, FPD), where lower distance reflects better coverage:

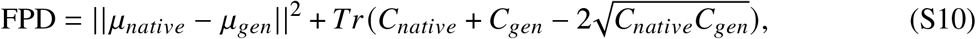

where *μ* is the feature-wise mean for each set of sequences, *C* is the respective covariance matrix, and *Tr* refers to the trace linear algebra operation, defined as the sum of the elements along the main diagonal of a square matrix. To infer function from a structural perspective, ESMFold was used to predict the structures corresponding to sequences generated by PROPEND. We then leveraged Foldseek to search the Protein Data Bank for proteins with the most similar structures in nature. A generation was considered “functional” if the probability of finding a similar protein structure was greater than 0.95 and the protein was annotated with the function described in the prompt. Each generated sequence was also evaluated for its similarity to native functional proteins. We used HHblits to search for homologs within the training dataset. For each of the alignments found in training dataset, we depict the one with the highest identity.

#### Generating protein sequences with the prompt of multi-objective

To redesign SagA, PROPEND and ProteinMPNN each produced 1,000 SagA sequences. To standardize the conditions, a temperature setting of 0.1 was maintained for both models, affecting sequence variability (Fig. S5). The sequences generated by PROPEND exhibited a range of identities with the wild type from 0.34 to 0.52, and the lowest sequence consistency observed among PROPEND’s outputs was 0.36, indicating a broad diversity spectrum. In contrast, ProteinMPNN generated 1000 SagA sequences with wild type identities ranging from 0.50 to 0.65, and the lowest sequence consistency among its outputs was 0.73, substantially higher than that observed with PROPEND.

### In vitro evaluation methods

#### TdT generation

TdT has an additional 13 kDa N-terminal region, which contains a nuclear localization sequence (*36*) and a protein-protein interaction BRCT-like domain (*37*) (Fig. 4B). Structures predicted by AlphaFold 2 revealed that the N-terminal region was predicted to contain highly flexible loops with low pLDDT, which might impact the correct folding of TdT when expressed in *E. coli*. It has been shown that removement of the N-terminal region has the potential to increase the expression level and activity of TdT (*38*).

#### SagA generation

The wild-type SagA gene was from Enterococcus faecium strains Com15 (Gen-bank WP_127808574.1). Full-length SagA contains a signal sequence (1-27 aa), N-terminal coiledcoil (31-268 aa), NlpC/P60 catalytic domain (389-530 aa) at C-terminal, and the N-terminus and NlpC/P60 domain was linked by a serine/threonine rich linker. Since N-terminus has been shown to inhibit the NlpC/P60 domain activity and only the 3-D structure of NlpC/P60 domain was available (*39*), we generated the novel sequences only for the NlpC/P60 domain with its structure (6B8C.pdb) as the input.

#### Materials

All reagents were purchased from Sigma-Aldrich unless otherwise noted. Synthetic genes were purchased from TsingKe Biotech, Beijing, China.

#### Expression and purification of TdT and SagA

The synthetic TdT genes were cloned into *BamHI* and *HindIII* on pET28a vector. BL21(DE3) strain was used for transformation with the plasmids encoding the TdT. The cells were grown in LB medium with 50 *μ*g/mL kanamycin and the cell culture were induced with the addition of isopropyl-*β*-D-thiogalactopyranoside (IPTG) with the concentration of 0.5 mM at 18 °C for 16 h after the OD600 reached 0.6. Cells were collected by centrifugation at 4000 rpm for 10 min and resuspended in 10 mL lysis buffer (50 mM potassium phosphate buffer, pH 7.5, 100 mM NaCl). The cells were sonicated for lysis and the cell debris were removed by centrifugation at 12000 rpm for 15 min at 4 °C. The protein was purified using Ni–NTA resin (Sangon Biotech, China). The supernatant was loaded onto a nickel affinity column under gravity flow. The columns were washed three times with washing buffer (50 mM potassium phosphate buffer, pH 7.5, 100 mM NaCl, 50 mM imidazole) in order to remove non-target proteins, then target proteins were collected with 10 mL elution buffer(50 mM potassium phosphate buffer, pH 7.5, 100 mM NaCl, 250 mM imidazole). The purified protein was concentrated by ultrafiltration using 10-kDa Centrifugal Filters (Millipore, Billerica, USA) with lysis buffer. The protein concentration was determined using Quick Start™ Bradford Protein Assay with BSA as the standard. The proteins purified were verified by Sodium dodecyl-sulfate polyacrylamide gel electrophoresis (SDS-PAGE).

#### Expression and purification of SagA

The synthetic SagA genes were coloned into pET28a vector encoding SagA N terminus, and expressed in BL21(DE3) strain. Transformed cells were grown for 12 h in LB medium supplemented with kanamycin. Cells were inoculated at 1:100 ratio in 50 mL fresh LB medium, grown at 37 °C, 220 rpm till OD600 reaching 0.6 and then induced by 1 mM IPTG for an additional 16 h at 18 °C. Cells were harvested by centrifugation at 4000 rpm for 10 min and resuspended in 15 mL lysis buffer (20 mM Tris-HCl pH 8.0, 300 mM NaCl, 5 mM imidazole). The cells were sonicated for lysis and then centrifuged at 12000 rpm for 20 min to remove cell debris. The protein in the supernatant was purified using a 1 mL nickel affinity column under gravity flow. The columns were washed with 20 mL washing buffer (20 mM Tris-Hcl buffer, pH 8.0, 300 mM NaCl, 20 mM imidazole) in order to remove non-target proteins, then target proteins were collected with 1 mL elution buffer(20 mM Tris-Hcl, pH 8.0, 300 mM NaCl, 500 mM imidazole). The purified protein was desalted using 10-kDa cut-off Centrifugal Filters (Millipore, Billerica, USA) with lysis buffer. The protein concentration was determined using Bradford Protein Assay. The proteins purified were verified by Sodium dodecyl-sulfate polyacrylamide gel electrophoresis (SDS-PAGE).

#### Enzyme activity measurement of TdT

The catalytic activities of TdTs were measured by the extension rate of a 23 nt primer with dideoxyguanosine triphosphate (ddGTP). ddGTP will terminate elongation of the primer due to the lack of a 3’ hydroxy group. The reaction system comprised 1 *μ*M oligonucleotides (primer N), 33 *μ*M ddGTP, 0.25 mM CoCl_2_, 50 mM phosphate buffer(pH 7.5) and 100 mM NaCl, 0.02 mg/mL purified protein. After incubation for 10 or 30 min at 37 °C, the reaction was quenched by heating at 98 °C for 2 min. Products were analyzed by polyacrylamide gel electrophoresis on 20% acrylamide gel and 8 M urea and run at 180 V for 80 min, then the gel was stained with NA-Red for 10 min and imaged on SHST Analysis (SHENHUA SCIENCE TECHNOLOGY CO., LTD). The primer N used is as Fig. S1 and N+1 was supposed to be the reaction product.

#### Enzyme activity measurement of SagA

The synthetic SagA genes were coloned into pET28a vector encoding SagA signal sequence and N terminus, and expressed in BL21(DE3) strain. The enzyme activities were measured by detecting the OD600 of cells expressing full-length SagA, as the expression of SagA would lead to cell lysis and hence cause a decrease in culture optical density (*40*). Transformed cells were grown for 12 h in LB medium supplemented with kanamycin. Cells were inoculated at 1:100 ratio in 800 *μ*L fresh LB medium in 96 deep-wells plate, grown at 37 °C, 220 rpm till OD600 reaching 0.2 and then induced by 1 mM IPTG. The cells without induction were also cultivated as control. The OD600 of the cell culture were measured at every 2 h and used for preparing growth curve. The activities of mutants relative to the wild type was calculated using following equation:

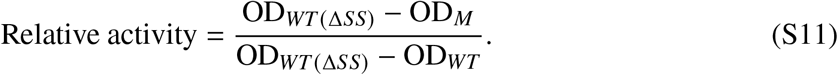

- OD_*WT* (Δ*SS*)_: OD600 of cells expressing SagA lack of signal sequence after cultivation for 8 h.
- OD_*M*_ : OD600 of cells expressing full-length SagA mutants after cultivation for 8 h.
- OD_*WT*_ : OD600 of cells expressing full-length wild-type SagA after cultivation for 8 h.

#### Melting temperature measurement of SagA

A nano differential scanning fluorimeter (nanoDSF) instrument, NanoTemper Technologies Prometheus NT.48 was used to determine the melting temperatures. SagA was always used at 500 µg/mL in the final solution. A temperature gradient of 1 °C/min from 20 °C to 90 °C was applied and the intrinsic protein fluorescence at 330 and 350 nm was recorded. The experiment was performed as three independent measurements.

**Figure S1:**
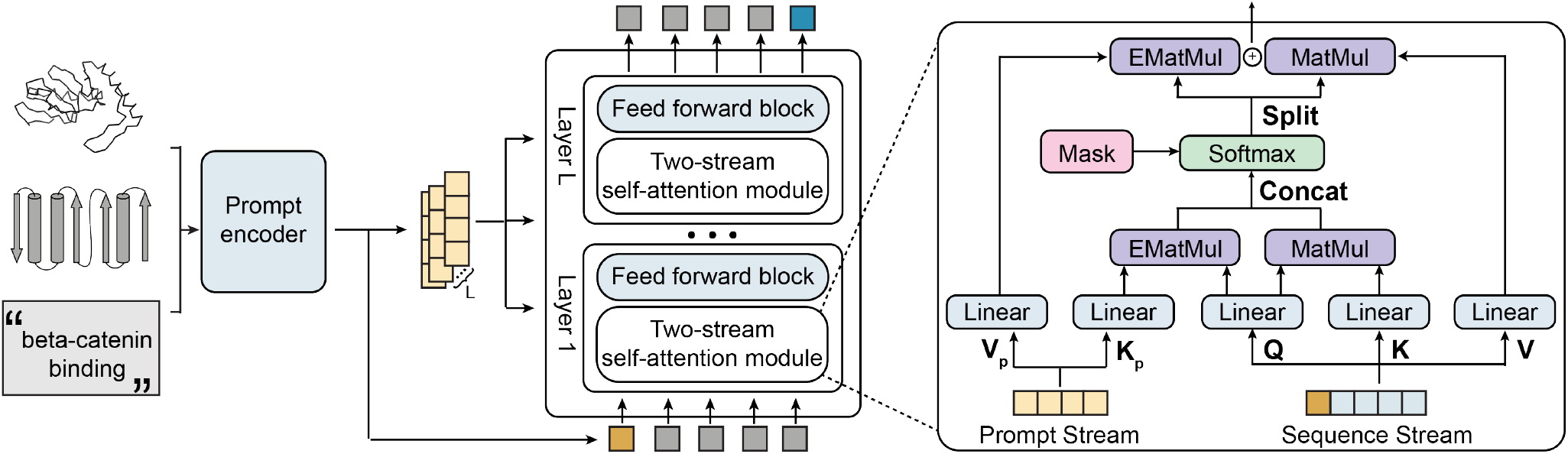
Model architecture. Arrows show the information flow among the various components described in this paper. The model is guided by the context and prompt to generate the next token. Element-wise matrix multiplication is used between the two stream to achieve a unique local prompt for each amino acid to be generated.

**Table S1:**
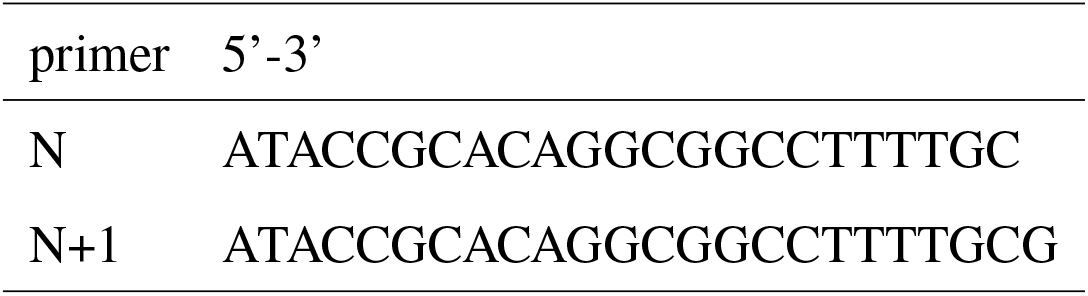
Primers used to measure TdT enzyme activity.

**Figure S2:**
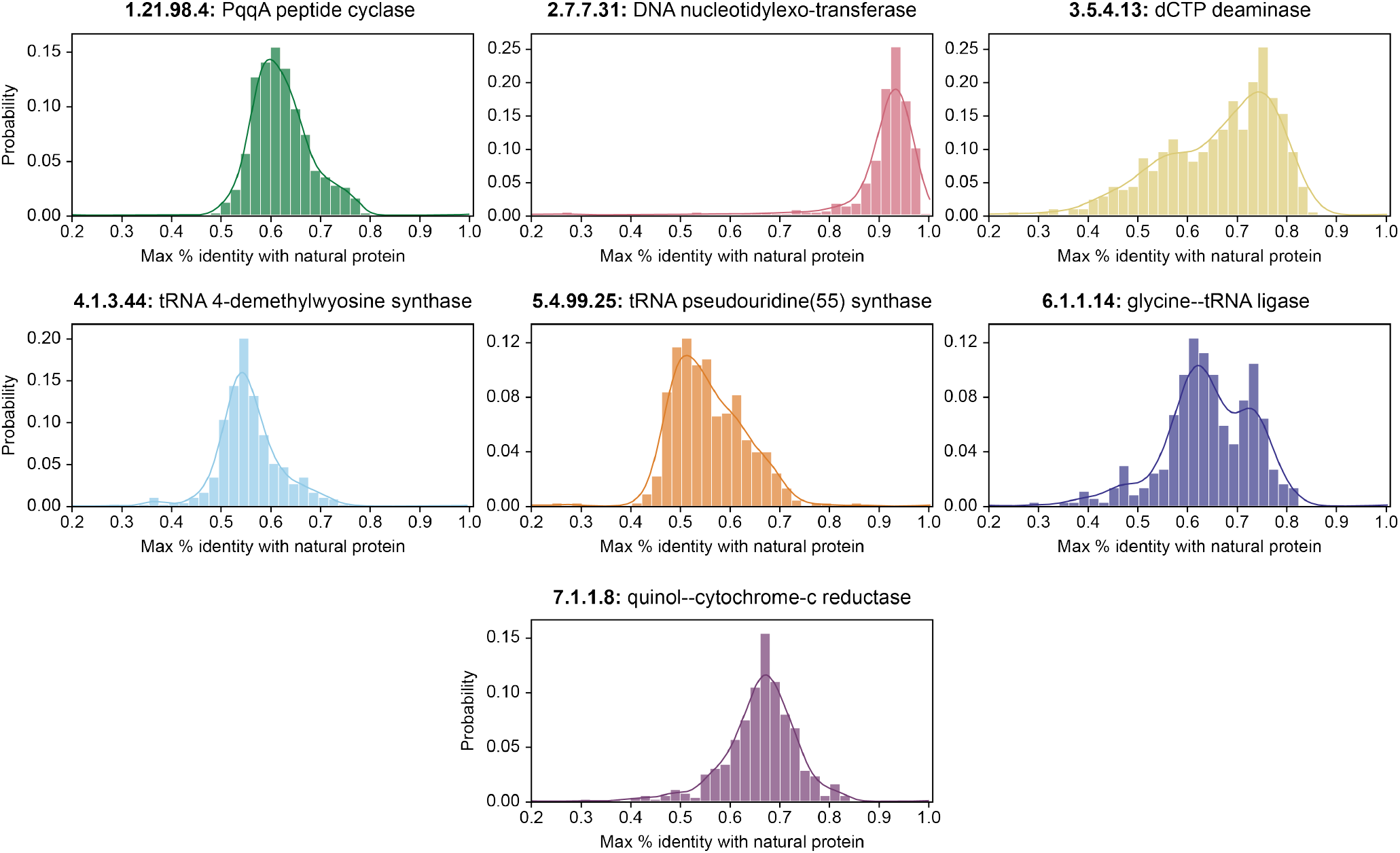
Distributions of sequence identity, over generated sequences from PROPEND using protein functions as prompts (n=500 sequences per condition).

**Figure S3:**
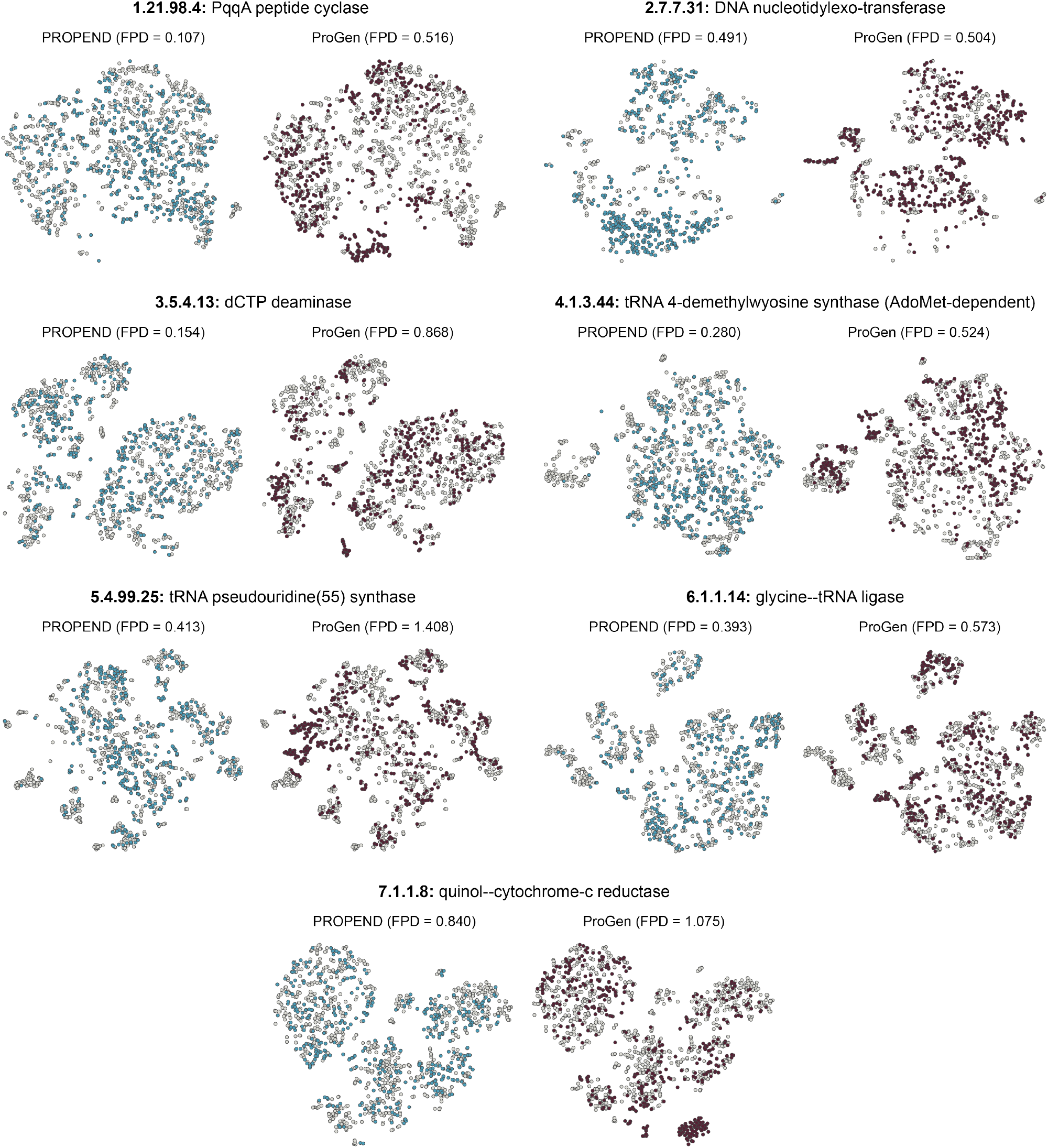
Exploration of the sequence and functional diversity in distributions produced by PROPEND and ProGen. This t-SNE visualization, based on ESM2 embeddings, compares natural sequences from the test set (represented in grey, n=1000) with those generated by PROPEND and a fine-tuned ProGen model (650M), shown in blue and dark red (n=500 for each), and is annotated with FPD values.

**Figure S4:**
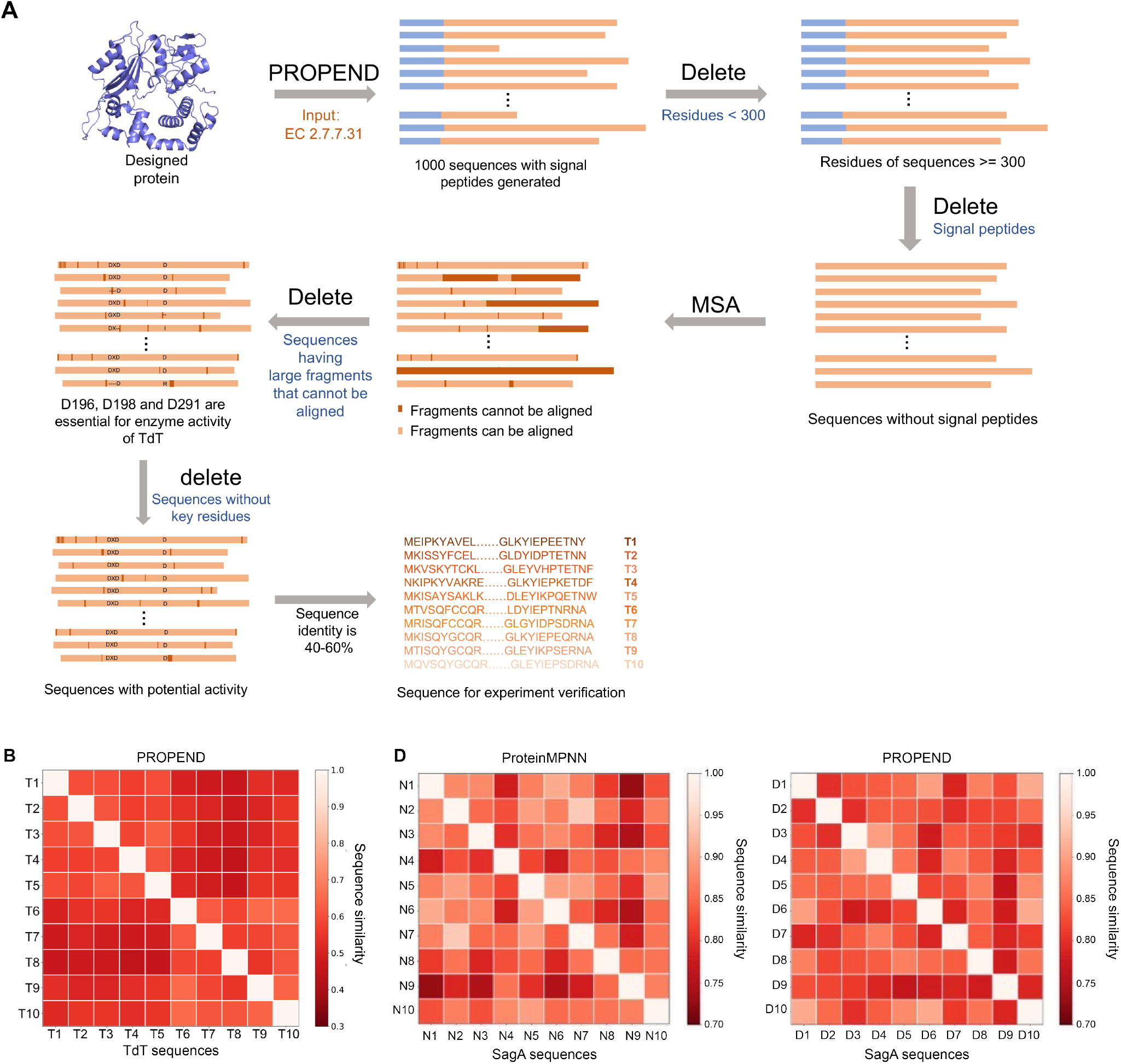
Screening process and sequence similarity of proteins verified by wet experiment. (**A**) TdT Generation and Selection Pipeline. Using “DNA nucleotidylexotransferase” as a prompt, PROPEND generated 1,000 sequences. We then refined this library based on sequence length, signal peptides, MSA alignment, key residues, and identity compared to the wildtype. (**B**) Similarity between generated TdT sequences from PROPEND. (**C**) Similarity between generated SagA sequences ProteinMPNN and PROPEND.

**Figure S5:**
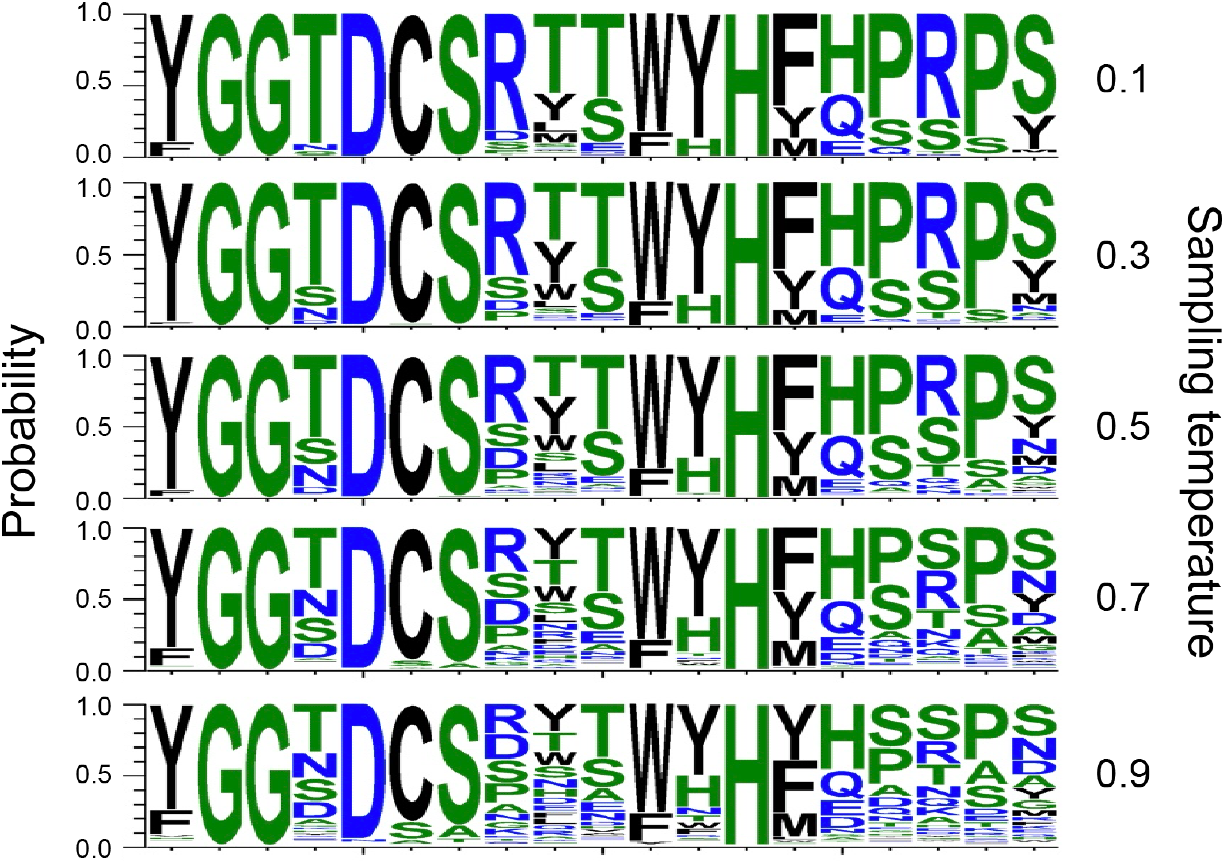
Relationship between the diversity of sequences generated by PROPEND and the sampling temperature.

